# eQTL network analysis reveals that regulatory genes are evolutionarily older and bearing more types of PTM sites in *Coprinopsis cinerea*

**DOI:** 10.1101/413062

**Authors:** Jinhui Chang, Tommy CH Au, CK Cheng, HS Kwan

## Abstract

Understanding the DNA variation in regulation of carbohydrate-active enzymes (CAZymes) is fundamental to the use of wood-decaying basidiomycetes in lignocellulose conversion into renewable energy. Our goal is to identify the regulators of lignocellulolytic enzymes in *Coprinopsis cinerea*, of which the genome harbors high number of Auxiliary Activities enzymes.

The DNA sequence of *C. cinerea* family including 46 single spore isolates (SSIs) from crosses of two homozygous strains are used to develop a panel of SNP markers. Then the RNA sequence were used to characterize the gene expression profiles. The RNA were extracted from cultures grown on softwood-enriched sawdust to induce lignocellulolytic enzymes and CCR de-repression genes. To assess the genetic contribution to enzyme expression variations among the 46 SSIs, associations between SNPs and gene expressions were examined genome-widely. 5148 local eQTLs and 7738 distant eQTLs were obtained. By analyzing these eQTLs, the potential regulatory factors of the CAZymes expression and the de-repression of Carbon Catabolism Repression (CCR) were identified,.

The eQTL network is characterized in terms of hotspots, evolutionary age and post-translational modifications (PTMs). In the eQTL network of *C. cinerea*, the non-regulatory genes are younger than the regulatory genes. The proteins regulated by combinational multiple types of PTMs are more likely to function as super regulatory hotspots in protein-protein interactions. The evolutionary age analysis and the PTMome analysis could serve as alternative methods to identify master regulators from genomic data.

This work demonstrates a comprehensive bioinformatics approach to identify regulatory factors with next-generation sequencing data. The results provide candidate genes for bioengineering to increase the enzyme production, which will practically benefit the bioethanol production from lignocellulose.

**Significance:** This eQTL analysis is designed to study the fungal CAZymes and carbon catabolism repression, especially during the mycelium stage.

1. In *Coprinopsis cinerea*, only the regions near two ends of the chromosomes have high recombination rate, and suitable for family based eQTL analysis.
2. A sugar transporter is a hotspot controlling many CCR genes.
3. CAZymes are not regulated by a master regulator, but by individual regulators. This indicates that CAZymes are under specific regulatory pathways, so can response to specific conditions.
4. In the eQTL network, the rGenes are evolutionarily older, with more types of PTM sites than eGenes.
5. In the eQTL network, the proteins with more types of PTM sites are more likely associated with Information Storage and Processing, and act as super-hub in the network.

## Introductions

Lignocellulose is the most abundant natural resource and its conversion into renewable energy attracts many research interests. Understanding the variation in regulation of carbohydrate-active enzymes (CAZymes) is fundamental to the use of wood-decaying basidiomycetes in lignocellulose conversion. Many of the CAZymes are important in biofuel production, waste processing and other industries, especially in lignocellulose degradation. Carbon catabolite repression (CCR) is the major repression mechanism controlling CAZymes expression, and leads to low production of enzymes industrially. “Bioprospecting” to uncover the regulating pathways of CAZymes will provide crucial information on their uses. Gene expression is a fine-tuned program involving signal transduction, chromatin conformational modification, transcription related protein recruiting and transcription, it is under regulation of various levels, from DNA, RNA to protein modifications. The focus of this study is to understand gene expression of enzymes most relevant to biomass degradation, at DNA level and protein level.

DNA variants among individuals alter the expression periods and levels of genes. Genetic loci associated with various gene expression patterns are defined as expression quantitative trait loci (eQTLs). eQTLs are divided into two categories according to the distance between the loci and the associated gene. If the eQTL is located around the gene they affect, it is called cis-eQTL or local-eQTL. If the eQTL regulate distant genes, it is called trans-eQTL or distant-eQTL.

Post-translational modifications (PTMs) are known to diversify the protein functions and dynamically modulate the signal transduction, hence regulate gene expression in various manners. On individual proteins, multiple types of PTMs may cooperate and interact. Interactions between PTM types have been reported in many proteins, and are proposed to be a universal mode of protein regulation. It is found that PTMs usually cooperate with each other either in positive or negative manner, by which generate robust dynamic responses (Hunter 2007). Theoretically, to discriminate a wide range of signals and to response accordingly seems hardly relies on limited number of signaling transduction cascades; while combinatorial use of PTMs on signal transducers or regulators, especially the key ones, seems like a necessary strategy.

eQTLs network can give more insight about how genetic variants regulate gene expression and perhaps how they are involved in complex phenotypes. The proteins will be characterized to examine if proteins with mtp-PTM are more likely as a hub in eQTL network, and if functional enrichment exists for mtp-PTM proteins.

*Coprinopsis cinerea* is a saprotroph generally growing on the non-woody substrates dung, while its genome harbors a high number of Auxiliary Activities enzymes. Through these analysis, a better understanding of the eQTL network will be obtained, and the regulatory mechanism of combinational interaction among multiple PTMs.

## Results and Discussions

### SNPs with high recombination rate are near the ends of chromosome

SNPs with low minor allele frequency (MAF) are non-informative (too little variation) and have a great potential for creating spurious findings. SNPs whose either genotype occurs fewer than 5 times (MAF of <10%) or have more than 10 missing data, were filtered out. Second, the genotypes of were compared with the genotypes of their parental strains. The parental strains are not perfect homozygous, and there are few of SNPs have more than two genotypes. Given the significant difference of the minor allele frequency (MAF) of the homozygous parental SNPs from that of the non-homozygous SNPs, the data with more than two genotypes were excluded for downstream analysis. There are 265,306 SNPs passed these filters.

The 265,306 SNPs were then plotted along the chromosomes (Figure 1. shows Chromosome 2, other chromosomes in supplementary figures). There are 46 layers representing the 46 SSIs. Green dots represent the genotype same as the reference genome and red dots for the genotype same as the mapping partner. Grey dots are missing data. White blank means that there is no difference between the reference and the mapping partner. Most proportions in the middle in each chromosome show similar pattern, indicating the recombination rate is low. There are tiny differences inside the long identical blocks, indicating double-crossovers occur frequently. Only the SNPs near two ends of the chromosomes have high recombination rate, and suitable for family based eQTL analysis.

**Figure 1.**
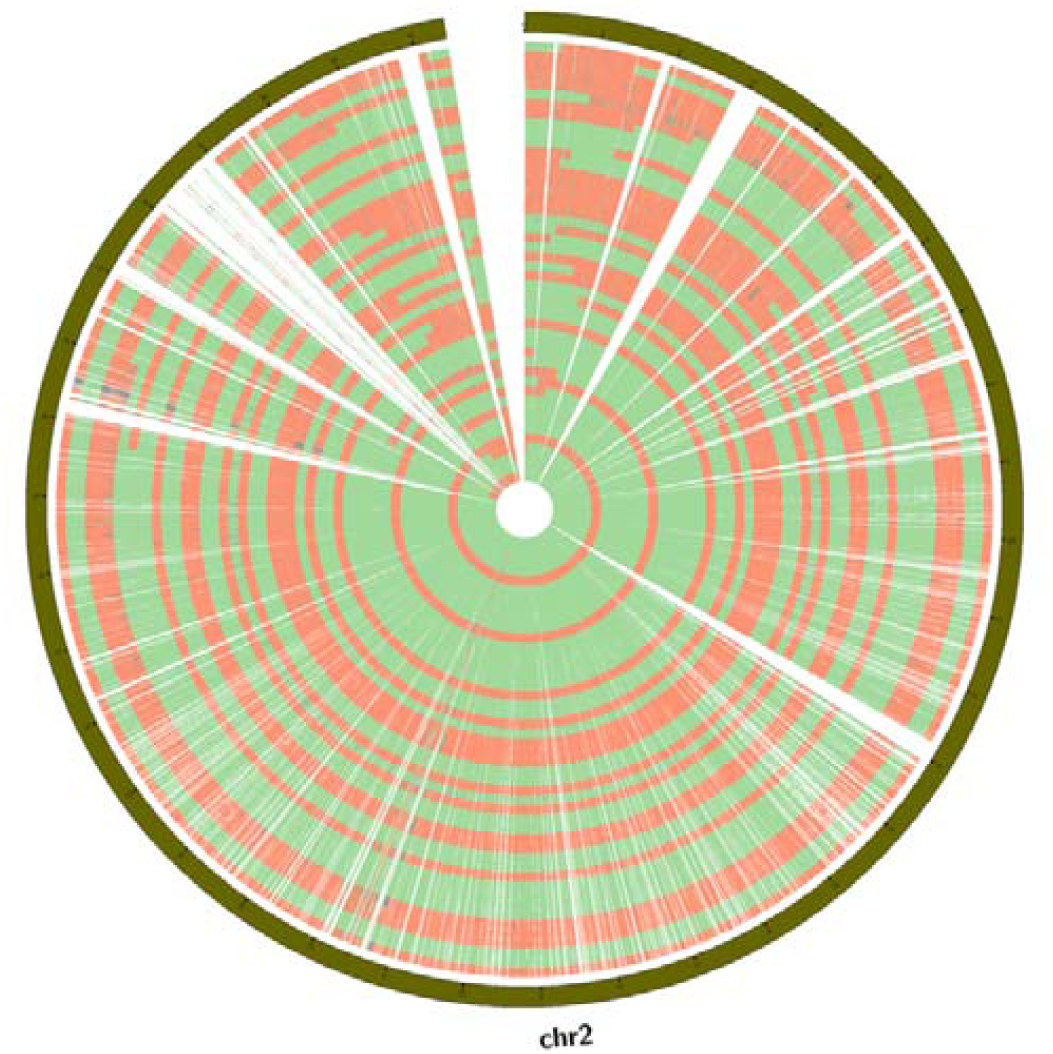
Genotype matrix of chromosome 2 in the 46 SSIs. There are 46 layers from inside to outside for each of the 46 SSIs. Green dots represent the genotype same as the reference genome and red dots for the genotype same as the mapping partner. Grey dots are missing data. White blank means that there is no difference between the reference and the mapping partner. The most outside layer shows the length, and the unit is 1 × 10^*5*^ *bp.*

### eQTL

The eQTL analysis for all gene expression and SNP genotype pairs are test for association with a *P-*value of no more than 1 × 10^−6^, and FDR rate no more than 0.001. The distance cutoff of 5 × 10^5^ bp between the gene and the SNP is set, to differentitate local and distant eQTLs.

The number of eQTL pairs passed *P*-value and FDR threshold is too large to handle. There are 4,727,512 pairs of gene expression and SNP genotype passed the thresholds for local eQTL association. The resulting local eQTLs have *P-*values from 1.6 × 10^−32^ to 9.9 × 10^−7^, FDR from 1.4 × 10^−24^ to 1.8 × 10^−5^. There are another 4,802,760 pairs passed the thresholds for distant eQTL association. The resulting distant eQTLs have *P-*values from 8.9 × 10^−32^ to 9.9 × 10^−7^, FDR from 3.1 × 10^−22^ to 7.2 × 10^−4^.

Since the recombination rate is low in the middle of the chromosomes, the SNPs along very long regions have same/similar genotype variation patterns, as shown in Figure 1. The very long regions contain too many genes and SNPs, and it is very difficult to locate the real causative ones. Therefore, these noisy long regions are not informative and should be discarded.

The count of eSNPs of each eGenes is used to screen the informative eQTLs. The cutoff is 100 for the number of eSNPs per eGenes. After this screening, there are 5148 local eQTLs and 7738 distant eQTLs left. These eQTLs are named as independent eQTLs.

### eQTL network and the regulatory hotspots

The rGenes and eGenes generated from the eQTL analysis can be used to build a gene-gene interaction network. Genes are the nodes, and the eQTL associations are the links, which is directed from the rGene to its corresponding eGene. Since there are many SNPs in one single rGene, the links between a pair of rGene and eGene can be repeated. There are also many self-loop in cases that the rGene and eGene are the same one. Most of genes are linked by the genes who can act as both eGenes and rGenes and interacted into a big network. There are also some small independent networks.

The eQTL hotspots are believed to regulate the expressions of a batch of genes, in manner of transcription factor, or upstream players of signal transduction pathways, or modifiers of chromatin conformation. Super regulatory hotspots in such network are potentially master regulators of various pathways. I use the NetworkAnalyzer (Assenov *et al.,* 2008) to calculate the Out-degree of each node (a gene in the network), after removing isolated nodes, small network components, duplicated links and self-loops. The rGenes with Out-degree =1 are defined as eQTL ordinary regulators, the rGenes with Out-degree =2 are defined as eQTL regulatory hotspots, and the rGenes with Out-degree >=3 are defined as eQTL super regulatory hotspots.

The top super regulatory hotspots are listed in Table 1. These top hotspots have outdegree higher than 10, which means that they have more than 10 eGenes whose gene expression levels are associated with them. Among the eGenes regulated by these top super regulatory hotspots, there are 6 cazymes which are highlighted in red color. Most CAZymes are not under centralized regulation from same regulators, indicating that the CAZymes have different regulation pathways and are expressed under different conditions.

CC1G_13691 is a sugar transporter which involves in CCR. It has effect both to local eGenes and distant eGenes.

Many top super regulatory hotspots are poorly characterized, indicates that they are evolutionarily more recent, and need to be further studied for their functions in regulating gene expression in response to nutrients depletion.There is no significant chromosomal enrichment of hotspots, while each chromosome has a few hotspots.

### Evolutionary analysis of the eQTL network

The genes are grouped as non-regulatory genes (OutDegree =0) and regulatory genes (OutDegree >=1). The quantile of the 2 groups are plotted in Figure 2. The mean of phylostratum of non-regulatory genes is significant larger than that of regulatory genes (5.209249 and 4.110485, p-value < 2.2e-16, Welch Two Sample t-test). This means that the non-regulatory genes are evolutionarily younger than regulatory genes.

**Figure 2:**
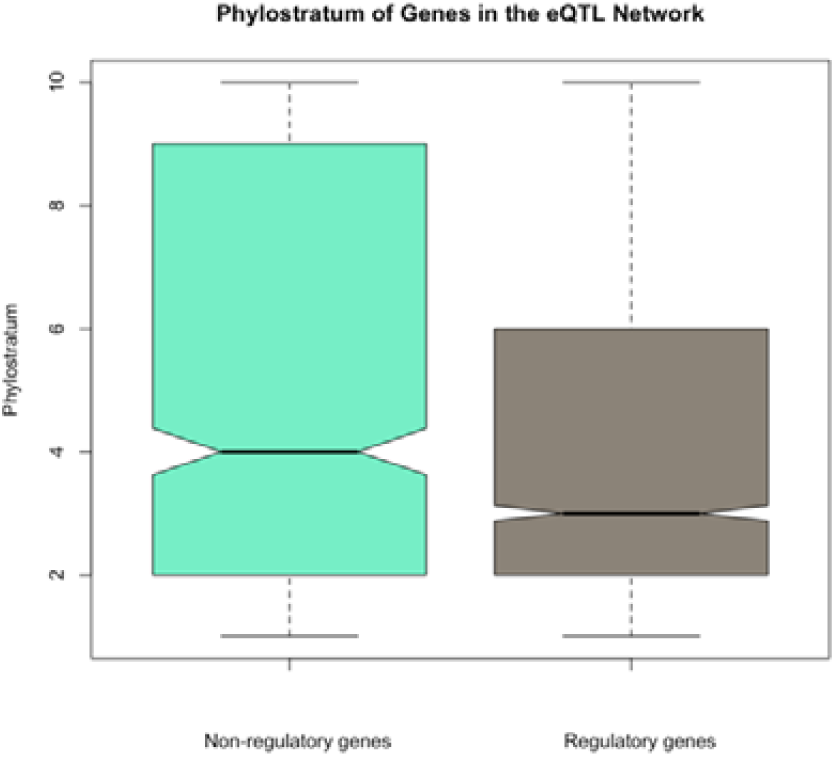
The non-regulatory genes are evolutionarily younger than regulatory genes. The lines in each box-plot represent the minimum value, 25%, median, 75% and maximum value, from bottom to top.

### *C. cinerea* Mtp-proteins enriched in regulatory hotspots in eQTL networks

Using the bioinformatic tools listed in supplementary methods section, the potential modification sites for 11 types of PTMs were predicted. If the proteins have 1 or 2 types of predicted PTM sites, it belongs to Group “<=2”. The proteins have 7 to 11 types of predicted PTM sites, are combined into Group “>=7”. The group sizes are 1775, 2277, 2634, 2652, 2063 and 1941, respectively.

Welch Two Sample t-test is used to compare the number of PTM types between eGenes and rGerens, and the results (t = 8.6738, df = 3066.2, p-value < 2.2e-16) support the alternative hypothesis that true difference in means is not equal to 0 at 95 percent confidence. The means of PTM types number is 4.322452 for eGenes and 4.824223 for rGenes. As shown in Figure 3a, rGenes have higher number of PTM types.

**Figure 3:**
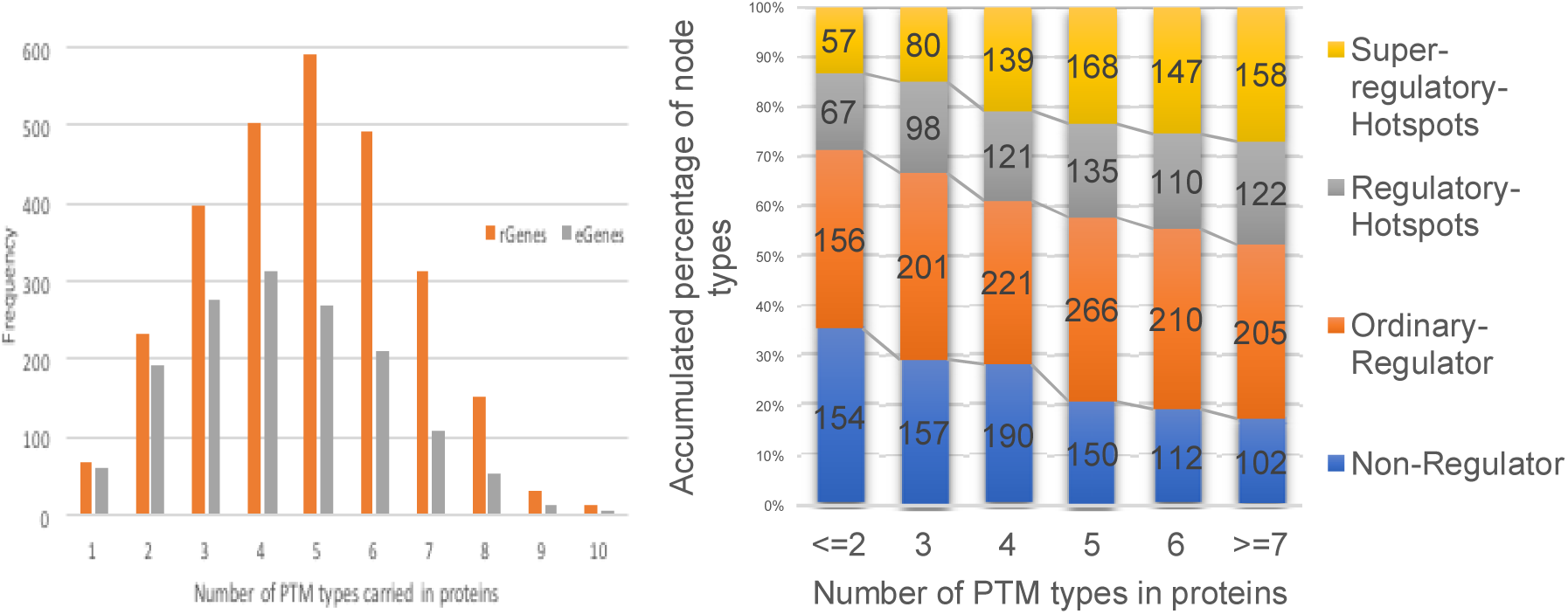
(a) Number of PTM types carried in eQTL proteins. *The count of rGenes and eGenes are presented by columns height. The x-axis is the number of PTM types carried in each protein.* (b) The accumulated percentage of node categories in different protein groups. *In the constructed eQTL network, all nodes were classified into four categories (Non-Regulator: OutDegree = 0; Ordinary-Regulator: OutDdegree = 1; Regulatory Hotspots: OutDegree = 2; and Super-regulatory-Hotspots: OutDegree* ≥ *3). The y axis shows the corresponding accumulated percentage of the different node types. This figure demonstrates that different protein groups present divergent roles in eQTL network.*

In the eQTL network, all nodes were classified into four categories (Non-Regulator: OutDegree = 0; Ordinary-Regulator: OutDdegree = 1; Regulatory Hotspots: OutDegree = 2; and Super-regulatory-Hotspots: OutDegree ≥ 3). Figure 3b demonstrates that different protein groups present divergent roles in eQTL network. Proteins harbouring more types of PTMs have more proportions to act as regulatory hotspots and super regulatory hotspots.

### KOG enrichment of independent eGenes

To characterize the independent eGenes, especially to find out what kind of genes are prone to be regulated by eQTLs, the functional enrichment of eGenes was examined. The eGenes of the independent eQTLs are grouped by KOG sub-class, and the count of genes in each group is shown in Figure 4 as columns. The percentage of eGenes in whole genome KOG-annotated genes in each KOG sub-class are also calculated and shown as line in the figure. The total number of eGenes is 8.9% of all KOG-annotated genes in the *C. cinerea* genome.

**Figure 4:**
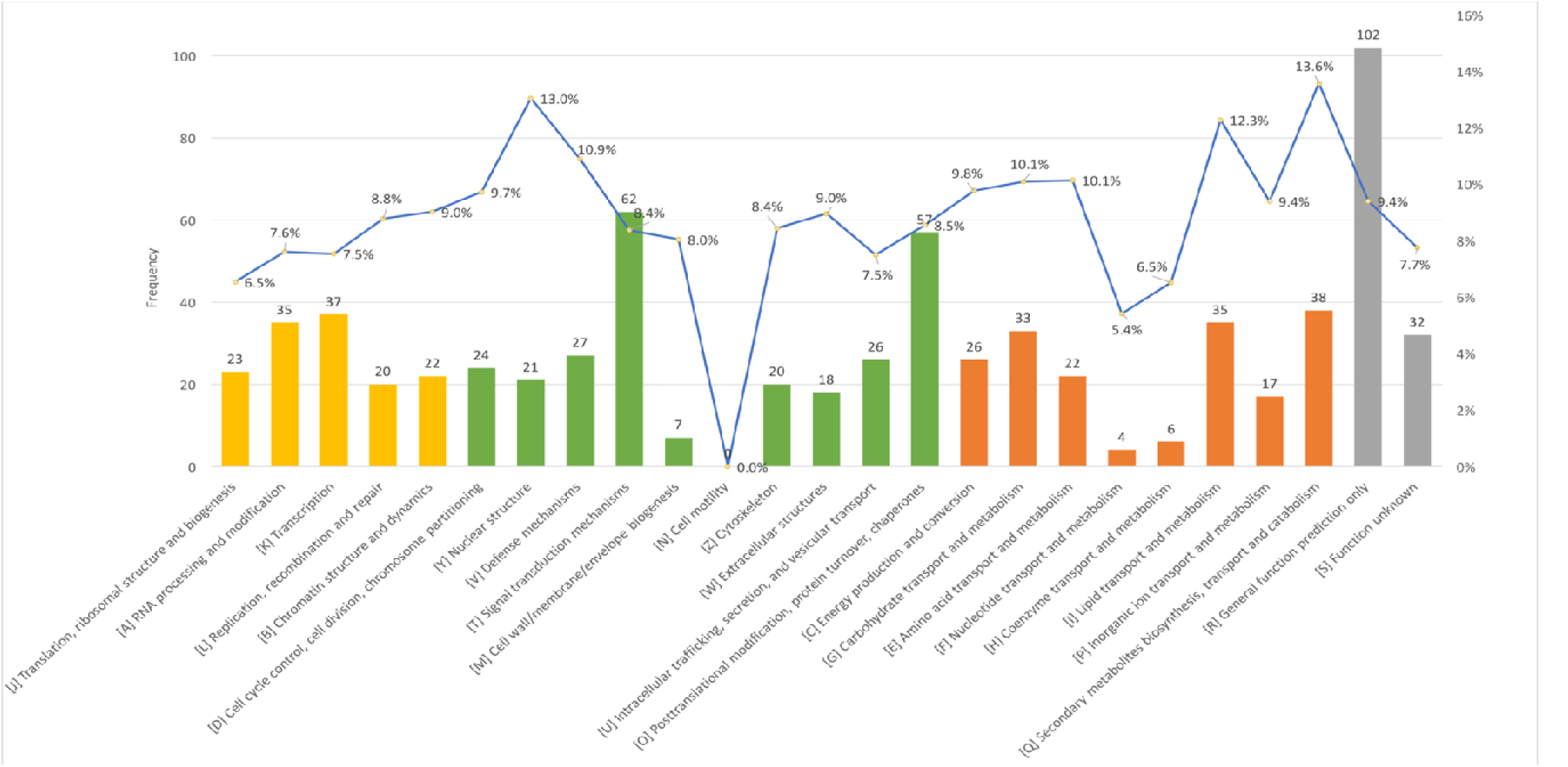
KOG enrichment of independent eGenes. Yellow bars are “INFORMATION STORAGE AND PROCESSING”, green bars are “CELLULAR PROCESSES AND SIGNALING”, orange bars are “METABOLISM”, and grey bars are “POORLY CHARACTERIZED”. The percentage of eGenes in whole genome KOG-annotated genes in each KOG sub-class are also calculated and shown as line in the figure.

The metabolism enzymes and structural components are prone to be eGenes. For example, 13.6% of “[Q] Secondary metabolites biosynthesis, transport and catabolism”, 13.0% of “[Y] Nuclear structure”, and 12.3% of “[I] Lipid transport and metabolism” are eGenes, which are much higher than 8.9%.

House-keeping genes have less chance to be regulated by eQTL, for example the genes from KOG sub-classes of [F] Nucleotide transport and metabolism, [H] Coenzyme transport and metabolism, [J] Translation, ribosomal structure and biogenesis, [U] Intracellular trafficking, secretion, and vesicular transport, [K] Transcription, and [A] RNA processing and modification, have lower percentage than 8.9% of eGenes. KOG sub-class “[N] Cell motility” which has only 4 members in the *C. cinerea* genome, and none of the 4 is a eGene.

### eQTL of CAZymes indicate there is no master regulator

Oxidases and carbohydrate-active enzymes related to lignin, cellulose and related aromatic compound degradation are readily retrievable from the CAZy (Lombard et al. 2014) database. To assess the genetic contribution to CAZymes transcriptional variation among 46 SSIs, eQTL analysis was performed for the 491 CAZymes.

Among the 181 rGenes, there is no master regulator who can regulate the expression of dozens of cazymes, while 31 rGenes can regulate 2-4 cazymes (Figure 5). 62 out of the 491 cazymes are independently under regulation of one or several rGenes. Local regulatory polymorphisms identified eQTL for the 62 CAZymes genes at 5% FDR. rGenes are potential regulators of CAZyme expression.

**Figure 5:**
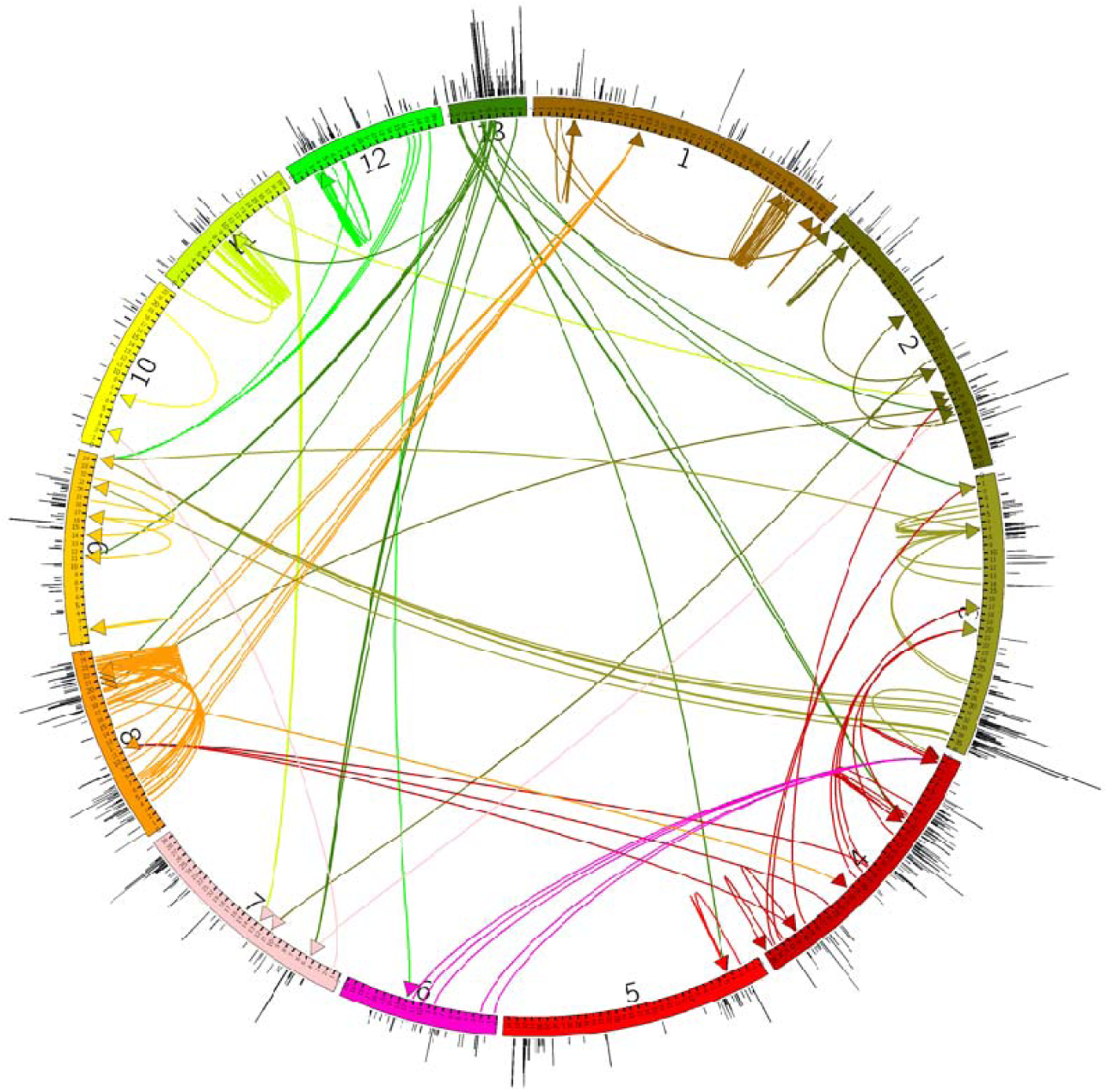
The positions of CAZymes and the corresponding eQTLs. The histogram outside the circle indicates the position of hotspots(Outdegree>=2). Color of the link is same with its rGenes’ chromosomes. Links showing the relationship and locations of the rGenes and the eGenes in each eQTL pair. Arrows point to the eGenes, which are CAZymes in this figure.

CAZymes are regulated by specific regulators, so the different enzymes are expressed in response to different conditions. This specificity is in accord with the finding in laccases. There are 17 laccases in C. cinerea genome, while only Lcc1 and Lcc5 are the main laccaes secreted in liquid cultures(Rühl et al. 2013).

Among the rGenes, there are 2 CCR components, 7 kinases and 5 transcription factors. Most microbes preferentially consume energy-efficient carbon sources first when several carbon sources are available. CCR is an essential regulatory system to enable microorganisms to uptake carbon sources efficiently, and balance the metabolic capacity (Wu et al. 2016). The rGene CC1G_00662, is one gene involved in the CCR. It is a Guanine nucleotide binding protein and a regulatory associated protein of mTOR, which usually involves in TOR signaling cascade, cell communication, cellular response to stimulus. The rGene CC1G_02575 is another CCR related gene. It is Cyclophilin type peptidyl-prolyl cis-trans isomerase B. It has cis-trans isomerase activity and involves peptidyl-amino acid modification, and thus cellular protein metabolic process.

When take a closer look at the links around the rGenes, some cases are highly likely to be causative genes of the CAZymes differential expression. One example is the pathway from sugar transporter, to transcription factor, then to an AA3_2 enzyme (Figure 6a). CC1G_13691 is a sugar transporter. The 2 SNPs on this gene has 12 eGenes and is a regulatory-hotspot in the eQTL network of this study. One of the eGenes is CC1G_10373, which is a Zn-finger protein. The SNP on CC1G_10373 has association with the expression level of CC1G_01540, an AA3_2 enzyme, which is a pyranose dehydrogenase and exhibits extremely broad substrate tolerance with variable regioselectivity (C-3, C-2 or C-3 + C-2 or C-3 + C-4) for (di)oxidation of different sugars. Although the function of CC1G_10373 is unclear, it could act as a regulator due to the zinc finger motif. Zinc finger domains typically serve as interactors, binding DNA, RNA, proteins or small molecules(MacPherson et al. 2006; Laity et al. 2001).

**Figure 6.**
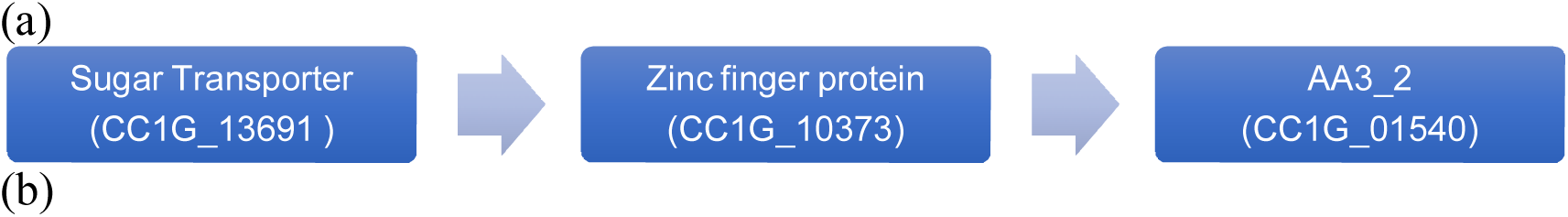
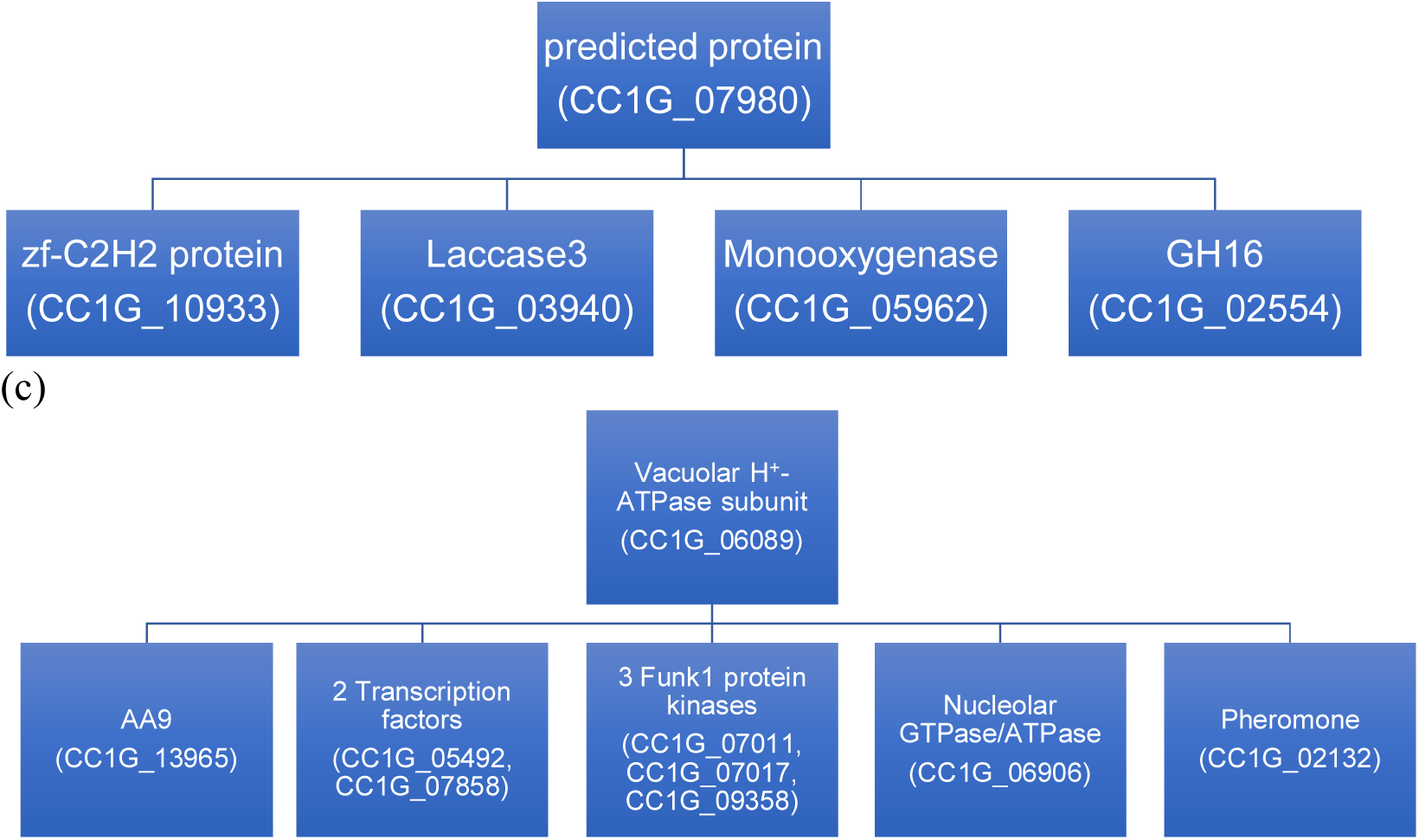
(a) The hypothetical regulatory pathway of CC1G_01540. (b) The eGenes of CC1G_07980. (c) The eGenes of CC1G_06089.

Another example of the hypothetical pathway is deducted from the eGenes of CC1G_07980, which is a predicted protein and poorly characterized. Its eGenes include: CC1G_10933, a zfC2H2 protein; CC1G_03940, laccase3, which is an AA1_1family multicopper oxidases; CC1G_05962, a Kynurenine 3-monooxygenase; CC1G_02554, which belongs to Glycoside Hydrolase Family 16. The homologue of CC1G_02554 in *Saccharomyces cerevisiae* is Crh2p/Utr2p, a glycosylphosphatidylinositol(GPI)-anchored chitin transglycosidase that transfers chitin from chitin synthase 3 to b-1,6-glucan, to create cross-links during cell wall biosynthesis. One possible illustration is the SNP on CC1G_07980, predicted protein controls the expression of CC1G_10933, which can bind to DNA and/or RNA and affect the expression of the laccase3, monooxygenase and the chitin transglycosidase (Figure 6b).

The last example of direct CAZyme regulatory pathway is deducted from the eGenes of CC1G_06089, a Vacuolar H+-ATPase subunit. Vacuoles are important as storage reservoirs for basic amino acids and polyphosphate, as sites for protein degradation and turnover, and as sequestration sites of potentially toxic ions, especially Ca2+. Vacuolar H^+^-ATPase can hydrolyze ATP and pump protons across membranes to acidify cellular compartments. The eGenes of CC1G_06089 include CC1G_13965, AA9(formerly GH61), copper-dependent lytic polysaccharide monooxygenases (LPMOs); CC1G_05492, a Homeobox transcription factor, which is a homologue of Sip1, and participate in Snf1-mediated CCR derepression in ascomycetes; CC1G_07858, HMG-box transcription factor; CC1G_06906, a nucleolar GTPase/ATPase; CC1G_02132, a pheromone protein (Phb2.2 B44); and three FunK1 protein kinases, CC1G_07011, CC1G_07017, and CC1G_09358. One possible illustration of this pathway is: the cell energy signal is sensed and transduced by the Vacuolar H^+^-ATPase subunit. The ATPase then pass the signal through FunK1 protein kinases cascades and affect the transcription of the two transcription factors, and hence the transcription of AA9 (Figure 6c).

### eQTLs of CCR-associated genes

Direct CAZyms eQTLs cannot explain all expression variations. Some cellulases and hemicellulases are clustered and transcribed together under complex regulation. Such regulations involve crosstalk of several signaling pathways, and Carbon catabolite repression (CCR) is one of the most important mechanism controlling the lignocellulolytic enzymes transcription. CCR enables microbes to utilize carbon sources efficiently, in which glucose represses the transcription of enzymes targeting less preferred carbon sources. The CCR-associated genes are predicted by blast and OrthoMCL. There are a lot of CCR components homologues in the eQTL, either as rGenes or eGenes.

Glucose sensing and the subsequent CCR are predominantly coordinated via the cAMP/protein kinase A (PKA) pathway in *S. cerevisiae*. There are two major routes of cAMP-dependent PKA activation, (1) extracellular glucose sensing via the Gpr1, G-coupled protein receptor and (2) the intracellular sensing of phosphorylated glucose (Figure 7a). G-protein alpha subunit homologue, CC1G_09275, and the hexokinase Hxk1 homologue, CC1G_11986, are both rGenes. The activation and localization of PKA also has cross-talk with TOR signaling. Sch9p plays a significant role in TOR-dependent nutrient sensing and is required for TOR-mediated ribosome biogenesis, entry into G0 phase, and has been shown to influence cell lifespan. The homologue of regulatory associated protein of mTOR, CC1G_00662, and the Sch9 homologue, CC1G_00081, are also rGenes. The Reg1p-Glc7p phosphatase dephosphorylate the activation loop of Snf1p inhibiting the use of alternative carbon sources. The Reg1 homologue, CC1G_11986, is a rGene.

**Figure 7:**
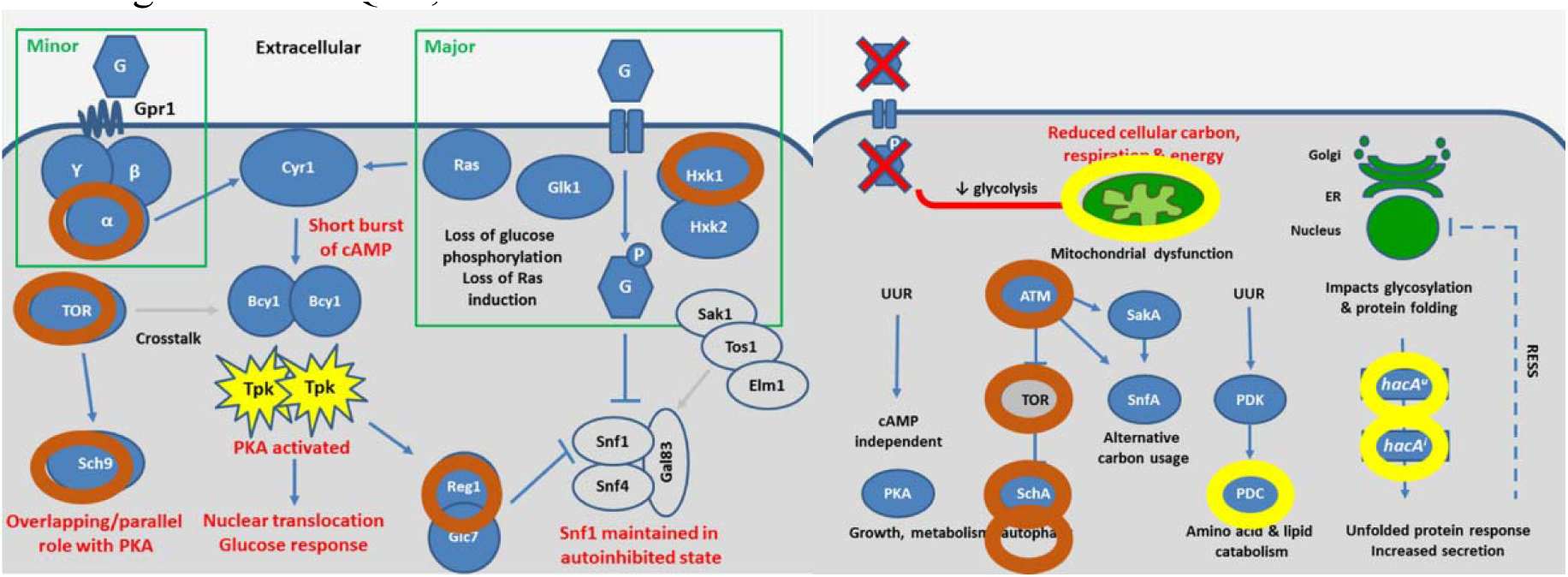
(a) The homologue of components of cAMP/PKA glucose sensing pathway in *S. cerevisiae* in the presence of glucose. (b) The homologue of components in the signaling pathways involved in CCR de-repression in *A. nidulans*. *The circle means that C. cinerea has the homologue of the circled gene. The red color means that the gene is rGene in this eQTL analysis, while the yellow color means that the gene is eGene in this eQTL analysis. Legend: grey = unactive; blue = activate; G = glucose; arrow = induction; bluntended arrow = repression; red X = absence of glucose; UUR = unidentified upstream regulator. Adapted from* (Brown et al. 2014).

There are also homologues of components in the signaling pathways involved in the carbon limitation stress response. Figure 7b shows the carbon limitation stress response activated in *Aspergillus nidulans* when lignocellulose is a sole carbon source. A reduction in cellular glucose causes a drop in intracellular energy levels and mitochondrial dysfunction. The ATM kinase senses mitochondrial dysfunction and activates alternative energy pathway including the TOR/SchA and SakA/SnfA pathways, and responses including autophagy, CCR derepression, and the secretion of hydrolytic enzymes. ATM homologue (CC1G_00839), TOR homologue (CC1G_00662), SchA (the Sch9p homologue in *A. nidulans*) homologue (CC1G_00081), and the autophagy Atg1p kinase homologue (CC1G_15592) are all rGenes. The mitochondrial ABC transporter homologue, CC1G_07837, is an eGene. The mitochondrial pyruvate dehydrogenase complex (PDC) represents an important metabolic switch for fuel selection, which during carbon starvation needs to be inactivated to switch carbon metabolism to fatty acid/amino acid catabolism. The homologue of PDC subunit alpha, CC1G_09082, is an eGene. The HAC1/hacA transcription factor induces the unfolded protein response (UPR). Triggering the UPR results in the up-regulation of the secretory capacity. The UPR and hacA splicing mechanism, is important to industry as a key mechanism to enable fungi to secrete sufficient lignocellulolytic enzymes under carbon limiting conditions. The hacA homologue, CC1G_05114, is an eGene.

### CCR-like systems in *C. cinerea*

The homologues of some CCR components of ascomycetes are absent in *C. cinerea*. There are no reports of CRE1 homologue in basidiomycetes, which is the master repressor of cellulases and hemicellulases genes in ascomycetes. Are there genes acting the roles of CRE1. As shown in Figure 8, the omponents of the CCR have three main roles: sensing, signaling and transcription regulation. It is suggested that the genes with these three functions as the main subjects to be investigated. Therefore, the focus were on the glucose/hexose transporters, G-proteins, ubiquitin ligases, phosphatases, kinases, transcription factors, chromatin binding proteins and histones in the eQTL results. Three CCR-like eQTL sub-networks were identified.

**Figure 8:**
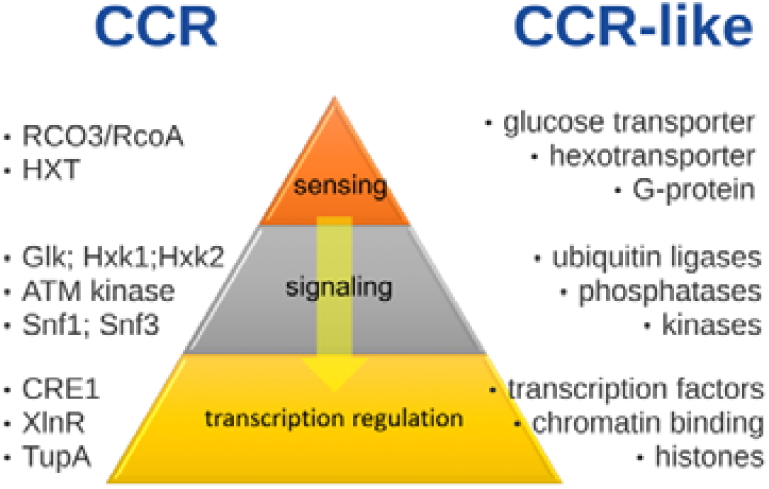
The comparison between CCR and CCR-like systems.

#### Sugar sensing and Transcription regulation

CC1G_13691 is a sugar transporter, and has 12 eGenes including 8 transcription related genes, one protease, cell cycle controller, ABC transporter and a cytoskeleton component. The WD40 repeat-containing protein CC1G_08510, is the TupA homologue, which is a corepressor of CCR.

#### Signaling and transcription regulator

CC1G_15592 is homologue of Atg1p, a highly conserved autophagy related gene. It is a kinase, responsible for the induction of autophagy. As explained in Figure 7b, autophagy will be turned on during starvation, while de-repression occurs. CC1G_11986 is a hexokinase, and a homologue of Hxk1p. It can phosphorylate the intracellular glucose and is involved the CCR induction, as shown in Figure 7a. CC1G_10330 is a serine/threonine protein kinase. CC1G_10329 is a polyadenylate-binding protein (RRM superfamily). The RNA binding zf-C3HC4 motif suggests it has transcription regulation capacity. These four rGenes have a common eGene, CC1G_10336, which is a transcription regulator DJ-1. This indicates a cross-talk among different signaling pathways will influence the expression level of a transcription regulator (Figure 9).

**Figure 9.**
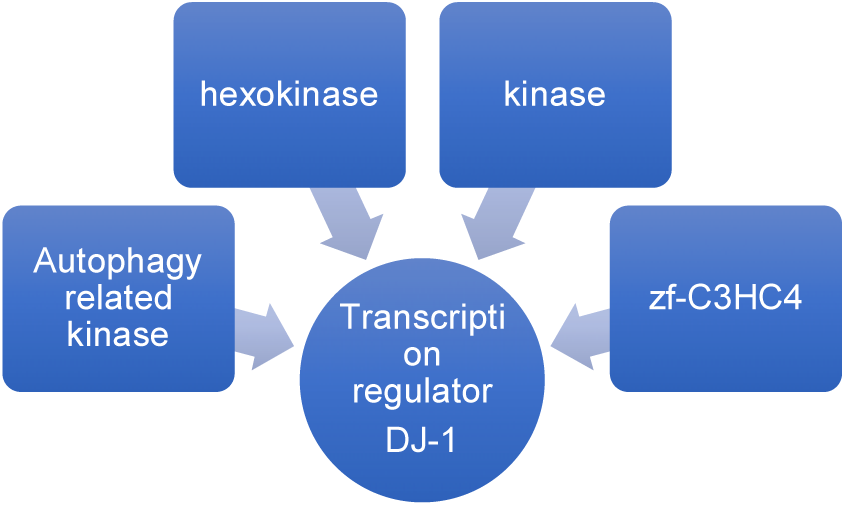
The signaling kinases and the transcription factor.

#### Singnaling (Repressor Degradation) and Transcription Regulation

During carbon starvation, the derepression will be induced via kinases/phosphatases signaling and repressor/TF degradation. CC1G_11801 is a F-box containing protein. The eGenes include pheromone/mating genes, transcription regulation related genes, lipid metabolism enzyme and a kinase.

### General characterization of the PTMome in *C. cinerea*

Ubiquitination is the most commonly known as the signal for protein degradation (Chen et al. 2015), and there are 5,946 proteins predicted containing more than one ubiquitination site. Methylation and Acetylation are primarily involved in epigenetics and chromatin state regulation (Shi et al. 2012). While more than half proteins have at least one acetylation sites, only 233 proteins are predicted having methylation sites.

Phosphorylation, the best-known and most common PTM with regulatory roles in many cellular processes such as developmental pathways and cell cycle (Xue et al. 2008). Since phosphorylation, by different families of kinases, is a very prevalent modification type, the software predicts that every protein in *C. cinerea* has at least one phosphorylation site. In other words, all proteins in *C. cinerea* can be modified by at least one type of PTMs.

### *C. cinerea* Mtp-proteins are Enriched in Information Storage and Processing

Besides the unannotated and poorly characterized proteins, there are 5498 proteins annotated with functional KOG terms and categorized into 23 KOG functional sub-classes. The number of proteins in each KOG sub-class are compared among the 6 groups. As shown in Figure 10, when the number of predicted PTM types increases, the count of proteins in “Information Storage and Processing” and “Cellular Processes and signaling” increases. This trend is also true even when the group sizes of group “6” and “>=7” are getting smaller. The most significant difference can be detected between group “<=2” and the other groups, which are proteins modified by multiple types of post-translational modifications (Mtp-proteins). It is clear to see from the comparative analysis that, the Mtp-proteins are enriched in all KOG sub-classes of the “Information Storage and Processing”, and most KOG sub-classes of “Cellular Processes and signaling” are modified by more PTM types. In other words, proteins involved in Information and Signaling are modified by more types of PTMs. In detail, the proteins having more types of PTM sites are enriched in the following KOG sub-classes:

**Figure 10.**
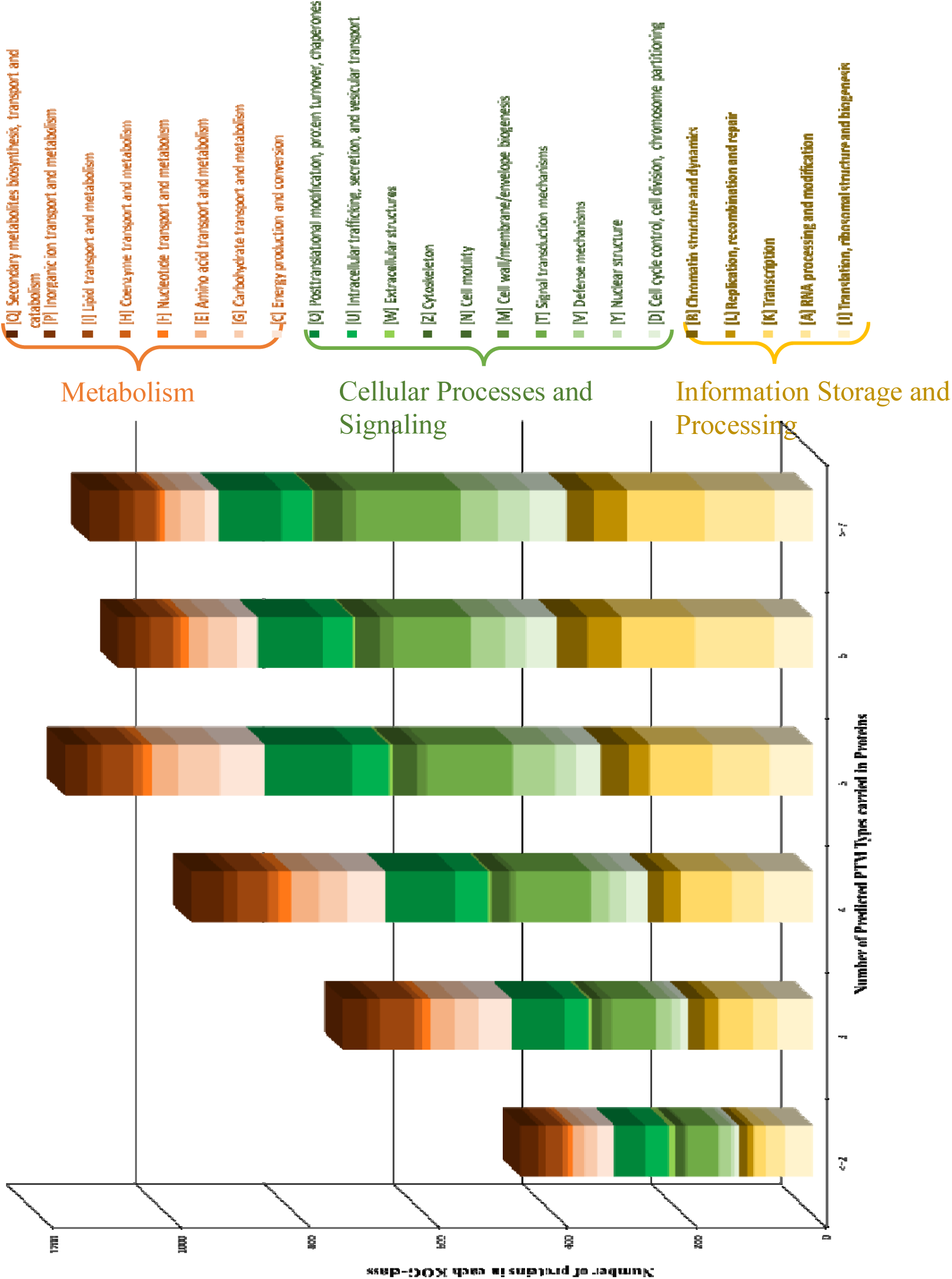
Number of Predicted PTM types v.s. KOG functional classes. The x-axis is the number of PTM types in proteins, and the y-axis is the count of proteins. The bands of different colors present the number of proteins in different KOG class. The yellowish bands present the KOG class of “Information Storage and Processing”; the greenish bands present the KOG class of “Cellular Processes and signaling”; the brownish bands present the the KOG class of “Metabolism”.

[J] Translation, ribosomal structure and biogenesis;

[A] RNA processing and modification;

[K] Transcription;

[L] Replication, recombination and repair;

[B] Chromatin structure and dynamics;

[D] Cell cycle control, cell division, chromosome partitioning;

[Y] Nuclear structure;

[V] Defense mechanisms;

[T] Signal transduction mechanisms;

[M] Cell wall/membrane/envelope biogenesis;

[N] Cell motility;

[Z] Cytoskeleton;

[U] Intracellular trafficking, secretion, and vesicular transport; and

[O] Posttranslational modification, protein turnover, chaperones.

At the same time, the count of proteins from the “Metabolism” category is getting smaller when the number of PTM types increases. In other words, the proteins involved in Metabolism are modified by fewer PTM types.

This enrichment agrees with comparative genomic analysis between mushroom-forming fungi and filamentous fungi. I compared the common KOG terms in mushroom-forming fungi and filamentous fungi. The genes in the group of “Posttranslational modification, protein turnover, chaperones” are expanded significantly. This KOG sub-class includes enzymes facilitate or catalyze the attachment or detachment of chemical groups to target proteins.

## Conclusions

New sequencing technologies have enabled high density SNP markers to be generated for mapping populations. This study demonstrates a way to investigate the genetic control of gene expression in mushroom forming fungi, while this method is suitable to be applied in all eukaryotic organisms.

### eQTL

eQTL mapping is a powerful tool to identify the genetic causes of gene expression variations, especially the SNPs correlate with lignocellulose degradation variation. A panel of 265,306 SNP markers were developed (∼1.1SNPs/KB). Among the 265,306 SNPs, 176,442 were mapped to the gene regions of 11629 genes, accounting for 87% of the total genes. On average, there are 15.2 SNPs per gene. SNP density in the genome is approximately one SNP per 139 □bp.

The straightforward but efficient and systematic eQTL method is used to identify regulators and genes under their regulation; there are 5148 local and 7738 distant independent eQTLs obtained. eGenes related to nutrient acquisition are enriched, particularly the Secondary metabolites biosynthesis, transport and catabolism, and lipid transport and metabolism, which points to the conversion of lipid derived acetate into energy rich carbohydrates under the experimental conditions.

In *C. cinerea*, the CCR mechanism is different from the one in yeast and filamentous fungi. The basidiomycetes do not have Cre1/Mig1 homologue, which can repress the cellulases and hemisellulases genes expression. The result of eQTL analysis shows that most rGenes can regulate only a few CAZymes. This may indicate that in this mushroom-forming fungus, which is more advanced than the yeast and filamentous fungi, the CAZymes are regulated in smaller scale.

This eQTL analysis also identify a series of CCR-like components, including sugar transpoter (sensor) and ubiquitin ligase (de-repression), which both have a large number of eGenes. Besides, transcription factors and kinases are also suggested as CAZyems regulators. They are good candidates of for future functional studies, and may be industrially important.

The eQTL network can also serve as a database to search gene-gene interactions, not only for CAZymes, but also for other functional groups of genes, as the sequence are from whole-genome DNA and RNA.

In the future, the functions of the predicted regulators should be further validated, and gene modified strains could be developed for industrial use. The results in this work will be of considerable practical importance to those interested in bioethanol production from cellulose-containing materials such as lignocellulose feedstocks, agricultural residues and sorted municipal wastes.

A limitation of our study is that the recombination rate is low and the sample size is small. The low recombination rate leads to the phenomenon that the genotypes of SNPs in a very long genomic region are too similar to discriminate the effect size of each SNPs, and made it impossible to locate the real causal SNPs. In the described work, the eGenes with more than 100 eSNPs were abandoned to solve this problem. However, this filtering results in some information lost.

The other limitation of the experimental design is that it is not designed in this project to distinguish between constitutive or induced responses to medium rich in lignocellulose for the mapping population, since I captured both responses by sampling mycelium under same culturing conditions, and did not replicate the whole trial with normal conditions.

I conclude that informative regulatory regions important to carbon metabolism can be identified by the association study of global gene expression and genome-wide genotyping data. Several these eQTLs can be shown to be specific to facilitate the lignocellulose utilization. This can help us understand the regulatory mechanisms underpinning genetic associations linked to CAZymes production.

This work characterized the eQTL network in terms of hotspot, evolutionary age and PTMs. The regulatory genes are evolutionarily earlier than the non-regulatory genes, and are more likely to be modified by multiple types of PTMs. These two features can be used to identify and/or validate the regulators in eukaryotes functional studies. This work not only expands the understanding of Mtp-proteins but also discloses the potential ability of Mtp-proteins to act as cores in protein-protein interaction.

The analysis reveals that Mtp-proteins are preferred to have more interactions in both mushroom forming-fungi. This finding can be supported by that many interactions are turned on/off in PTM dependent manner (Benkirane et al. 2010; Ho et al. 2008; Kruse & Gu 2008; Paolinelli et al. 2009). In cellular signaling network, multiple PTMs acting on one protein can enhance the effect and amplify the signal when needed. The regulation of recognizing or selectivity in PPI can be realized by creating or altering the docking sites for protein modules. Centrality-lethality rule holds that deletion of a hub protein has more lethal effect than deletion of a non-hub protein (He & Zhang 2006). So, the multiple modified proteins may play core roles in cellular biological processes and the proteins with no PTM site, locating in the periphery of the network, occupy less essentiality. It further implies the critical role of Mtp-Proteins in regulation of cell response.

### Insights in *C. cinerea* PTMome

PTM prediction is performed for all proteins in *C. cinerea*. The proteins are then characterized to examine if proteins with mtp-PTM are more likely as a hub in eQTL network. Then the enrichment and corresponding functions of combinatorial multiple types of PTMs, is analyzed.

PTM, especially the Mtp_PTMs, can be used to identify proteins which function as master regulators and superhubs. Since the PTM interact with other PTMs, it is a good mechanism to modulate its activity step by step. Mtp-proteins were found significantly enriched in protein complexes, having more protein partners and preferred to act as hubs/superhubs in eQTL network and protein-protein interaction (PPI) networks. They possess a distinct functional focus, such as chromatin assembly or disassembly, and reside in biased, multiple subcellular localizations. Mtp-proteins carrying PTM types biased toward locating in the ordered regions were mainly related to protein-DNA complex assembly.

The analysis of predicted PTM sites of the genome reveals that the proteins regulated by combinational multiple types of PTMs are tend to be important regulatory proteins. This provides us an approach to identify key components through *in silico* analysis of genome data.

One limitation of this study is that the PTM sites prediction is not accurate enough and have inflation. GPS 3.0 prediction results show that all 13442 proteins in C. cinerea have at least one phosphorylation site, of which many could be false-positive due to the algorithm specificity and accuracy limitation. However, this overestimated result agrees with the prevalence of protein phosphorylation, that more than 70% of all human proteins have been experimentally detected with more than one phosphorylation sites till the publication of (Olsen et al. 2010). Although I chose parameters for the highest accuracy in all predictors, the prediction results still be more or less overestimated.

Second, the sequences themselves of Mtp-Proteins also account for the observations. I found the length of protein is not the key point for multiple partners through correlation analysis. Although one protein can be extremely large and can have surface areas of thousands of Å^2^ units, it is received the restraints not only from physicochemical feasibility, but also from the cell and genome. It is impractical to increase binding sites simply by increasing the protein length.

Lignocellulose is the most abundant natural resource and its conversion into renewable energy attracts many research interests. Understanding the DNA variation in regulation of carbohydrate-active enzymes (CAZymes) is fundamental to the use of wood-decaying basidiomycetes in lignocellulose conversion. Our goal is to identify regulators of lignocellulolytic enzymes in *Coprinopsis cinerea*, of which the genome harbors high number of Auxiliary Activities enzymes. This study shows that, by analyzing the genome and transcriptome data of a well-designed population, key candidate regulators could be identified.

This work demonstrates a comprehensive bioinformatics approach to identify regulatory factors with next-generation sequencing data. The results of this thesis provide candidate genes for bioengineering to increase the enzyme production, which will practically benefit the bioethanol production from lignocellulose.

Through these analysis, a better understanding of the proteins regulated by combinational multiple PTMs is obtained, and the regulatory mechanism of combinational interaction among multiple PTMs.

## Materials and methods

### Strains

In this project a family based population was used to conduct the association analysis. The mapping population is 46 single spore isolates (SSIs) randomly sampled from the spores generated from crosses between the reference, sequenced strain (Okayama 7 #130) and the mapping partner used to define hot spots for meiotic recombination (#172). The genome resequencing of the 46 SSIs will enable SNP detection, and the RNA sequencing will enable gene expression level profiling, hence eQTL analysis of factors involved in biomass degradation.

### Culture conditions

The strain was cultivated on YMD medium containing 0.4 % yeast extract, 1% malt extract and 0.4% potato dextrose solidified with Bacto ^®^ agar. Mycelium was cultivated on agar plates at 37°C for about 1 week until the mycelium grew over the whole agar surface. The incubator was kept at a relative humidity higher than 60% for the development of fruiting bodies.

To prepare RNAs, the parental strains and 46 SSIs were grown on YMD agar for 1 week, then the mycelium from each plate were transferred into one metal filter, which allows the mycelium to have access to the medium outside the filter. The metal filters were placed on the medium (softwood-enriched sawdust) to maximize the induction of lignocellulolytic enzyme genes contributing to lignin degradation.

### DNA and RNA extraction and sequencing

The DNAs were extracted with DNeasy Plant Mini Kit (Qiagen) from two biological replicates for each segregant of mycelium. Total RNAs were extracted by the RNeasy Plant Mini Kit (Qiagen) from two biological replicates for each segregant of mycelium. The DNA and RNA sample were then shipped to DOE Joint Genome institute for sequencing (Project ID: CSP_633).

200ng of DNA/ cDNA of each segregant was sheared to 270bp using the covaris E210 (Covaris) and size selected using SPRI beads (Beckman Coulter). Using the KAPA-Illumina library creation kit (KAPA biosystems), the fragments were treated with end-repair, A-tailing, and ligation of Illumina compatible adapters (IDT, Inc). Illumina Hiseq was used to perform the 150bp paired end sequencing.

### Bioinformatics processing

The bioinformatics processing includes four major sections: 1. SNP calling from genome resequencing data; 2. Gene expression calling from transcriptome data; 3. eQTL mapping; 4. Analyzing the eQTL network.

### eQTL mapping

Matrix eQTL was used to identify eQTLs by performing a linear regression between expression levels and genotypes. Here, I assume that the genotypes have only additive effect on expression. Matrix eQTL can report the following information: Test statistic (t-statistic of T-test), Effect size estimation, p-value by linear regression, and False discovery rate (FDR) estimated with Benjamini–Hochberg procedure.

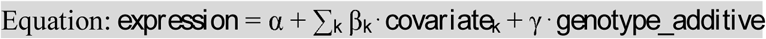

Outliers in expression data are transformed as the measurements for each gene into normally distributed while preserving relative rankings, with the R code provided in Matrix eQTL for such transformation:

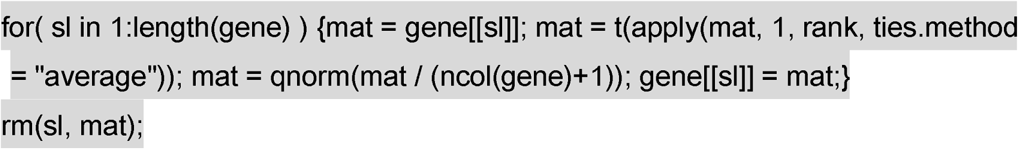

The local-eQTLs includes SNPs located within 0.5 Mb of the proximal gene. Correspondingly, associations between SNPs and genes outside this window are distant-eQTL. The eQTL analysis for all gene expression and SNP pairs are test for association with a *P* value of no more than 1 × 10^−6^, and FDR rate no more than 0.001. In each pair of eQTL, the SNP is denoted as eSNP, and the gene is denoted as eGene. The resulting eQTLs are then characterized by counting the number of affected eSNPs of each eGene associated with. The eGenes with fewer than 100 eSNPs are processed in the following analysis.

### Data analysis and functional enrichment analysis

All the SNPs are annotated by their locations. The eSNPs that can be mapped inside genes are denoted together with corresponding genes, which are named as rGenes. In this way, all the rGenes and eGenes can be used to build a gene-gene interaction network, using Cytoscape 3.4.0. And after removing isolated nodes, small network components, duplicated links and self-loops, the NetworkAnalyzer (Assenov et al. 2008) was utilized to calculate the Out-degree of each node (a protein in the network). The eSNPs with Out-degree >=3 are defined as eQTL hotspots in this work. Super-hotspots in such network are potentially master regulators of various pathways.

All the rGenes and cGenes are also annotated with KOG terms, KOG sub-class and KOG class, according to their functional domains. Functional enrichment analysis of the eGenes is performed to identify which group of genes have more chances to be regulated than by expectation.

### eQTL network construction and hotspots identification

The Cytoscape 3.4.0 is used to construct the eQTL network. Genes are the nodes, and the eQTL associations are the links, which is directed from the rGene to its corresponding eGene. I use the NetworkAnalyzer (Assenov *et al.,* 2008) to calculate the Out-degree of each node (a gene in the network), after removing isolated nodes, small network components, duplicated links and self-loops. The rGenes with Out-degree =1 are defined as eQTL ordinary regulators, the rGenes with Out-degree =2 are defined as eQTL regulatory hotspots, and the rGenes with Out-degree >=3 are defined as eQTL super regulatory hotspots.

### The evolutionary age of genes

The evolutionary age for each gene is estimated based on phylostratigraphy, which reveals evolutionary adaptation. It uncovers the footprints of important adaptive events in evolution. A protein is mapped to the time point—the phylostratum (PS)—when its oldest domain emerged in evolution. The *C. cinerea* genes are assigned to ten phylostrata in accordance with the NCBI phylogeny and the genome sequences available in the fungal taxa (Cheng et al. 2015). Smaller PS means evolutionarily older gene.

### Predictions and annotation of the PTMs

I use software to predict 11 types of PTMs, and the specific tools used for each PTM is listed in Table 3.2. GPS-series tools are using GPS (Group-based Prediction System) algorithm. The latest Linux versions of all programs were downloaded and installed on Centos5 64bit server. The parameters were set for highest accuracy.

### Classification of the proteins based on PTM pattern

Different proteins harbor the target residues of different number of PTM types, in other words, some proteins may carry the residues only targeted by one type of PTM and some others may contain the target residues of several PTM types. To disclose the potential special features and functions of proteins regulated by multiple PTM types (Mtp-Proteins), which at least have the target residues of two PTM types, I classified the proteins into different groups based on the number of PTM types carried in individual protein. Thus, the group “1” denotes that the proteins in this group only contain the target residues of one PTM type, and other groups by analogy.

### Functional characterization

To detect the potential function bias of Mtp-Proteins, I first assigned the functional annotations of different protein groups with KOG terms, KOG sub-class and KOG class. Various features of different groups were analyzed. Values in different groups were compared using the Welch Two Sample t-test.

